# Unveiling interactions of spatial-temporal information in tactile motion perception

**DOI:** 10.1101/2023.12.11.571089

**Authors:** Boyi Qu, Xiaojun Tan, Zheng Tang, Robert M. Friedman, Haiming Wang, Lan Lan, Kenneth E Schriver, Gang Pan, Hsin-Yi Lai

**Affiliations:** College of Biomedical Engineering and Instrument Science, Zhejiang University, Hangzhou, China; Department of Neurology of the Second Affiliated Hospital, Interdisciplinary Institute of Neuroscience and Technology, School of Brain Science and Brain Medicine, Zhejiang University School of Medicine, Hangzhou, China; Division of Neuroscience, Oregon National Primate Research Center, OHSU, Beaverton, USA; Department of Psychology and Behavior Science, Zhejiang University, Hangzhou, China; Liangzhu Laboratory, MOE Frontier Science Center for Brain Science and Brain-Machine Integration, State Key Laboratory of Brain-machine Intelligence, Zhejiang University, Hangzhou, China; College of Computer Science and Technology, Zhejiang University, Hangzhou, China; NHC and CAMS Key Laboratory of Medical Neurobiology, Zhejiang University, Hangzhou, China

## Abstract

The intricate interplay of spatial and temporal information in tactile-motion perception remains elusive. Despite strides in decoding neural signals for direction, speed and texture in tactile perception, nuanced interactions persist as challenges. Addressing this, we investigated direction and speed tactile perception, exploring the intricate spatial-temporal dynamics. Psychophysical experiments manipulated direction and speed parameters using a laboratory-designed fingerpad ball-stimulator. A pivotal discovery includes quadrant-dependent anisotropic distortion in perceived motion direction, expanding the well-known notion of a specific preferred orientation. Spatial features primarily influence inherent responses, while temporal features impact stimulus-specific responses, shedding light on dynamic directional perception. The introduction of a psychometric function improved the modeling of tactile-motion speed perception, incorporating both linear and nonlinear components for a more accurate representation. This study provides intriguing insights into the neural mechanisms in tactile-motion perception, with potential applications for somatosensation in brain-machine interfaces.

**Teaser:** This study unveils the intricate interplay of perceiving tactile motion, shedding light on the mysteries of tactile sensations.

## Introduction

In manipulating moving objects, precise tactile feedback at the point of skin contact is crucial for a secure grip (*1, 2*) and influencing object motion (*3-5*). Considering the example of catching a ball, where it is vital to judge its speed and location while adjusting grip strength and posture. Active touch enhances tactile information, contributing to improved perception of surface textures (*6-9*) or local contours (*10*). Thus, continuous interaction between dynamic tactile input (involving direction and speed) (*11*) and adaptable movements (*12*) provides a feedback loop for real-time acquisition of tactile spatial and temporal information (*13*). However, the integration of dynamic tactile information into our sensory experience remains poorly understood.

Skin mechanoreceptors play a pivotal role tactile-motion perception, with distinct properties influencing coherent motion perception by tracking spatiotemporal features (*9, 14-16*). Tactile inputs quickly update internal representations, enabling dynamic motion control (*2, 17*) and predictive strategies (*18, 19*). In somatosensory cortex, neurons exhibit robust direction selectivity (*20, 21*). Unlike mechanoreceptive afferents that relay information using temporal patterns (*20*). This difference in encoding could contribute to the pronounced perceptual biases in human tactile perceptions (*22*).

Tactile speed perception lags direction perception due to its intricate connection with spatiotemporal features, particularly in periphery texture modulation (*9, 23*). Recent studies highlight a complex interplay involving both speed-sensitive and non-sensitive neurons, challenging our understanding of neural encoding (*24*). Depeault et al. (*25*) emphasized the significance of spatial features in scaling perceived speed, showing a linear relationship with scanning speed. Recent research suggested complex interactions and nonlinear relationships in tactile sped perception, influenced by temporal features and local vibrations at different frequency (*26*). These findings underscore the roles of spatial and temporal features in tactile speed perception. However, the translation of dynamic features into generalized motion estimates in tactile perception remains unresolved.

This study investigates interactions between spatial and temporal features in tactile motion processing. Participants engaged in psychophysical experiments, perceiving the direction and speed of a tactile ball rotating on a distal fingerpad. We explored perceptual biases, directional patterns, and the potential relationship of spatial-temporal features and perceived speed. Analyzing the anisotropic pattern of perceptual bias in tactile motion direction, we systematically examined systematic and nonsystematic biases, revealing the potential role in integrating spatial and temporal information. Additionally, we assessed the impact of different stimulus parameters on perceived speed and compared mathematical models to understand the effects of stimulus speed and texture on tactile speed perception. Findings indicate that spatial and temporal features from external stimuli contribute to distinct aspects of tactile-motion perception, particularly regarding tactile speed. We used computational modeling to elucidate potential nonlinear mechanisms, thereby advancing our understanding of tactile-motion perception.

## Results

To explore perceptual bias and the integration of spatial-temporal information in tactile-motion perception, we designed an experimental setup to present a diverse of tactile-motion stimuli, encompassing various spatial and temporal features. These stimuli were presented on the left index fingerpad, and participants replicated the motion trajectory using their right index finger on a touchpad (Figure 1A). Our lab-designed stimulator, equipped with three degrees of freedom, enabled the investigation of motion stimuli with co-varied direction, speed, and wavelength, allowing us to explore the interaction of spatial and temporal features in tactile-motion perception (Figure 1B). We recruited thirty-three participants with normal tactile perception, and randomly divided into two groups. *Experiment I* focused on unraveling the perceptual bias symmetry and exploring the impact of speed on perceptual bias of a distal fingerpad (Figure 1C). In *Experiment II*, we delved into the impacts of spatial and temporal features on tactile-motion speed perception (Figure 1D). Additionally, mathematical approaches were employed to gain deep insights into the potential roles and interactions of these features in the tactile-motion perception.

**Fig. 1.**
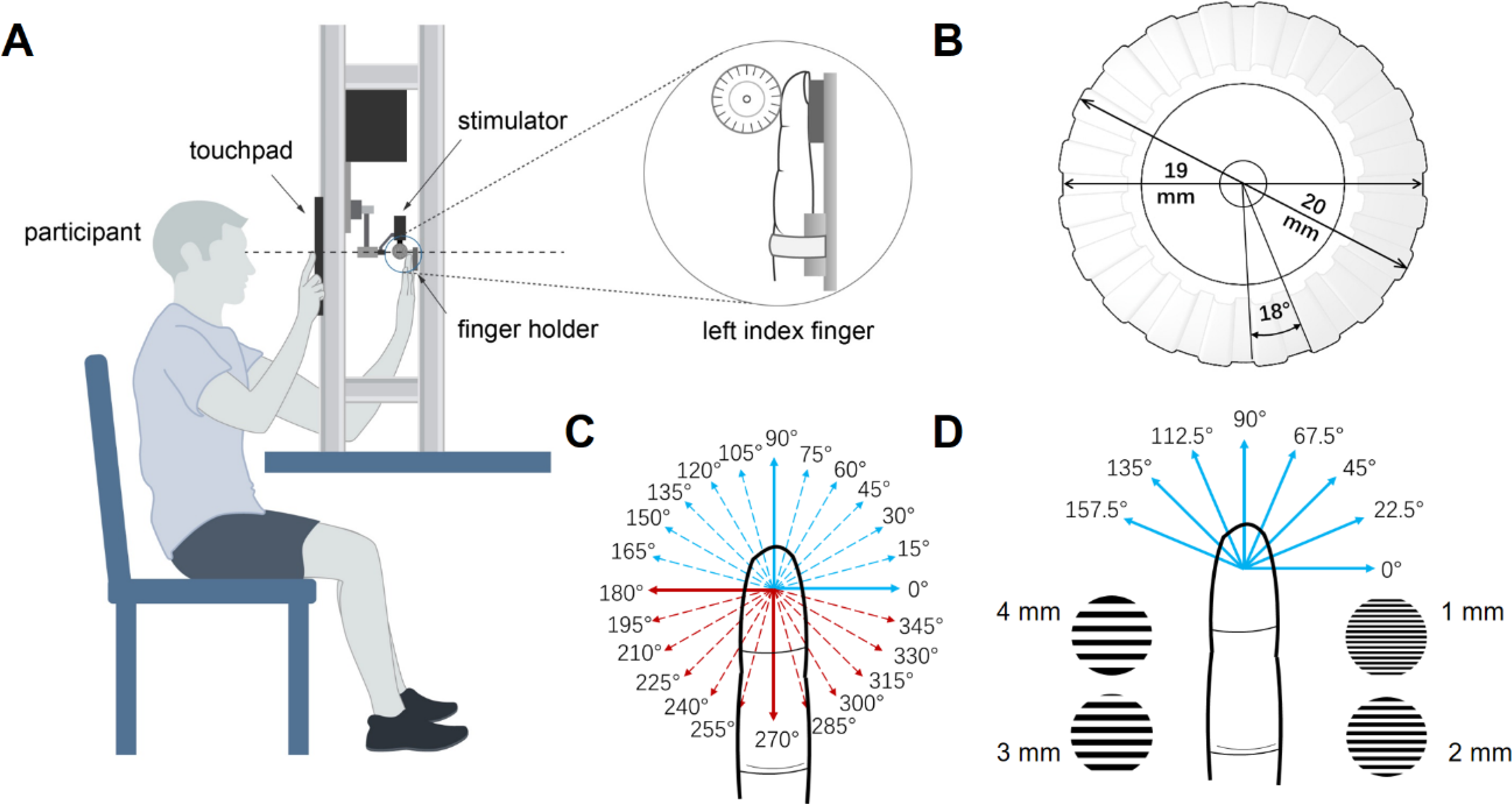
Setup and experiments design. (A) The participant is seated with their left index finger securely positioned, while using their right hand to indicate the perceived direction and speed of the tactile-motion stimulus on the touchpad. A computer controls the tactile stimulator and records data from the touchpad. The left index finger is held vertically within the apparatus, and the initial position of stimulus ball is set on the tangent plane between the fingerpad and the ball for each participant. (B) A schematic representation of a stimulus ball with a 3-mm wavelength is illustrated. The ball has a diameter of 20 mm and the grating indentation is 0.5 mm. The wavelength is defined as the maximum interval between the gratings, which are separated by an 18° step for a 3-mm wavelength ball. The angle of gratings is 5.62, 11.25, and 22.5 for 1-mm, 2-mm, and 4-mm wavelength balls, respectively. (C) Direction symmetry experiments utilized a comprehensive range of stimulus directions, spanning from 0° to 360° with 15° intervals. (D) In the tactile-motion speed integration experiment, eight motion directions were included, ranging from 0° to 157.5° with 22.5° intervals. Additionally, six distinct speeds (20, 40, 80, 160, 240, 320 mm/s) were used for each of the four wavelengths of the stimulus ball (1, 2, 3 and 4 mm).

### Anisotropic distortion in the perception of tactile-motion direction

*Experiment I* aimed to investigate perceptual bias symmetry of the distal fingerpad and the influence of speed on tactile-motion perception. The left index finger was stimulated with 24 directions under two speed conditions, covering the entire fingerpad. Traces drawn by right hand were pooled together following a feature extraction procedure (see Methods). Directional perceptual bias was calculated, and further analysis was conducted to evaluate the effect of speed and potential pattern indirection perception.

Perceptual bias, representing the deviation between the perceptual and veridical directions of the stimuli. We defined the mean perceptual bias, averaged across subjects, as systematic bias *B*_*s*_. In Figure 2A, the mean perceptual bias is depicted at the stimulus speeds of 40 mm/s (blue line) and 240 mm/s (red line) plotted against stimulus direction. Perceptual bias values consistently ranged from 0° to -45°, indicating a clockwise rotational shift. The bias tended to decrease as stimulus directions shifted from horizontal clockwise to vertical direction, exhibiting a known oblique effect (*27, 28*). Interestingly, increasing stimulus speed did not alter the overall anisotropy or systematic bias (*B*_*s*_ = -16.64° ± 0.82° SEM for 40 mm/s, *B*_*s*_ = -16.46° ± 1.02° for 240 mm/s; paired t-test: T (408) = -0.199, p = 0.842).

**Fig. 2.**
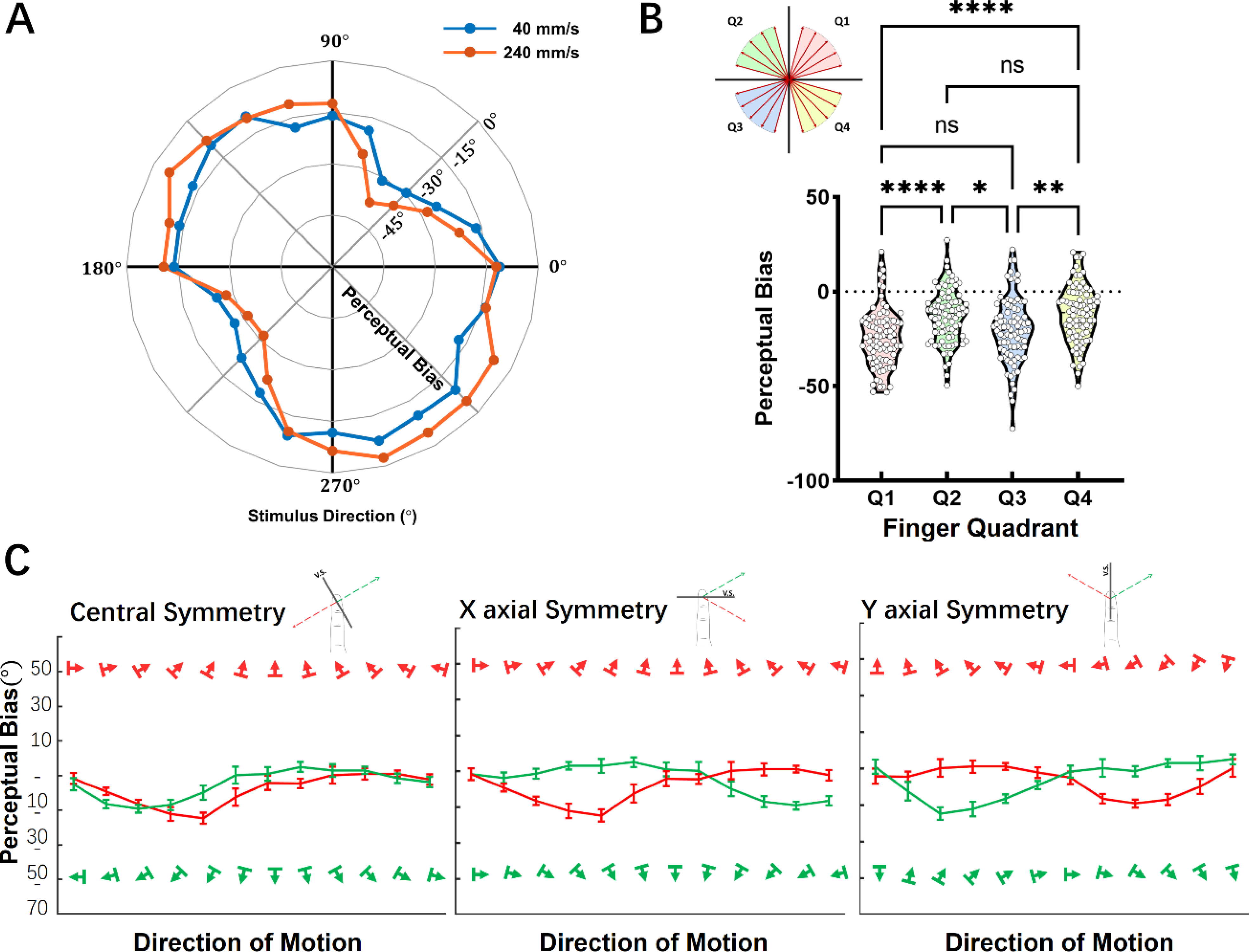
Anisotropic distortion in the perception of tactile-motion direction. (A) Perceptual biases are represented for the fingerpad with angles representing stimulus directions. No significant difference in systemic bias were observed between speeds of 40 mm/s and 240 mm/s. (B) Perceptual biases are divided into four quadrants (Q1, Q2, Q3, and Q4) for pairwise analysis. Quadrants in diagonal relationship exhibit noticeable similarities between them (Tukey, * represents p < 0.1, ** represents p < 0.01, **** represents p < 0.0001). (C) Direction perceptual bias is evaluated with respect to central, X-axial, and Y-axial symmetries. Colored markers indicating stimulus direction. A significant correlation is observed at the central symmetry, while no correlation is exhibited at X- and Y-axial symmetries. Data is presented as the mean ± the standard error of mean (SEM).

A trend was observed in Figure 2A, at higher speeds, there is an overall reduced perceptual bias in the second quadrant (Q2, upper left) and fourth quadrant (Q4, lower right), whereas there is an overall greater perceptual bias in the first quadrant (Q1, upper right) and third quadrant (Q3, lower left). Further analysis using one-way ANOVAs revealed significant modulation in the anisotropy pattern except for two diagonally related quadrants (Figure 2B, F(3, 340) = 12.69, p < 0.0001, with post-hoc in Tukey’s showed both non-significant results for Q1-Q3 and Q2-Q4). A stronger correlation was found between the perceptual bias in central symmetry distribution compared to X-axial and Y-axial symmetry distributions (Figure 2C, Pearson correlation, Left: Central symmetry, r = 0.679, p = 0.02; Middle: X-axial symmetry, r = 0.187, p = 0.58; Right: Y-axial symmetry, r = 0.156, p = 0.76). These findings support that the presence of anisotropies across the entire fingerpad, with a notable resemblance observed between the diagonal quadrants, supporting the existence of central symmetry during the direction perception. Moreover, the scanning speed does not significantly influence this anisotropy pattern.

### The role of speed and spatial frequency on tactile-motion direction perception

Investigating the influence of spatial and temporal features on tactile-motion speed perception, we used stimuli with varying spatial frequencies (ball wavelength), rotating at different speeds in *Experiment II*. To streamline conditions, we selectively stimulated a subset of motion directions covering the distal aspects of the fingerpad (Figure 1D) and four ball wavelength conditions, each presented in eight directions and six speeds. We introduced systematic bias and nonsystematic bias to assess dynamic representations, and mathematics models were employed to illuminate the speed perception.

Systematic biases at stimulus speeds of 20 mm/s, 40 mm/s, 80 mm/s, 160 mm/s, 240 mm/s, and 320 mm/s were -13.76°, -14.68°, -13.08°, -11.96°, -11.34° and -11.44°, respectively (represented by the different colored solid lines in Figure 3A that run parallel to the dashed zero bias line). Overall, systematic bias showed no significant difference between the stimulus speeds (F (5,384) = 0.824, p = 0.533) and averaged -12.71° ± 0.61°, indicating a clockwise shift as observed in *Experiment I*. Nonsystematic bias was fitted using a cosine function (as defined in Eq. 4) for various stimulus speeds. The peak amplitudes of the nonsystematic bias were 15.49, 15.85, 16.58, 18.42, 18.00, and 18.21°, while the corresponding phases were 0.366, 0.748, 1.09, 0.79, 0.963, and 1.251° for stimulus speeds ranging from 20 mm/s to 320 mm/s.

**Fig. 3.**
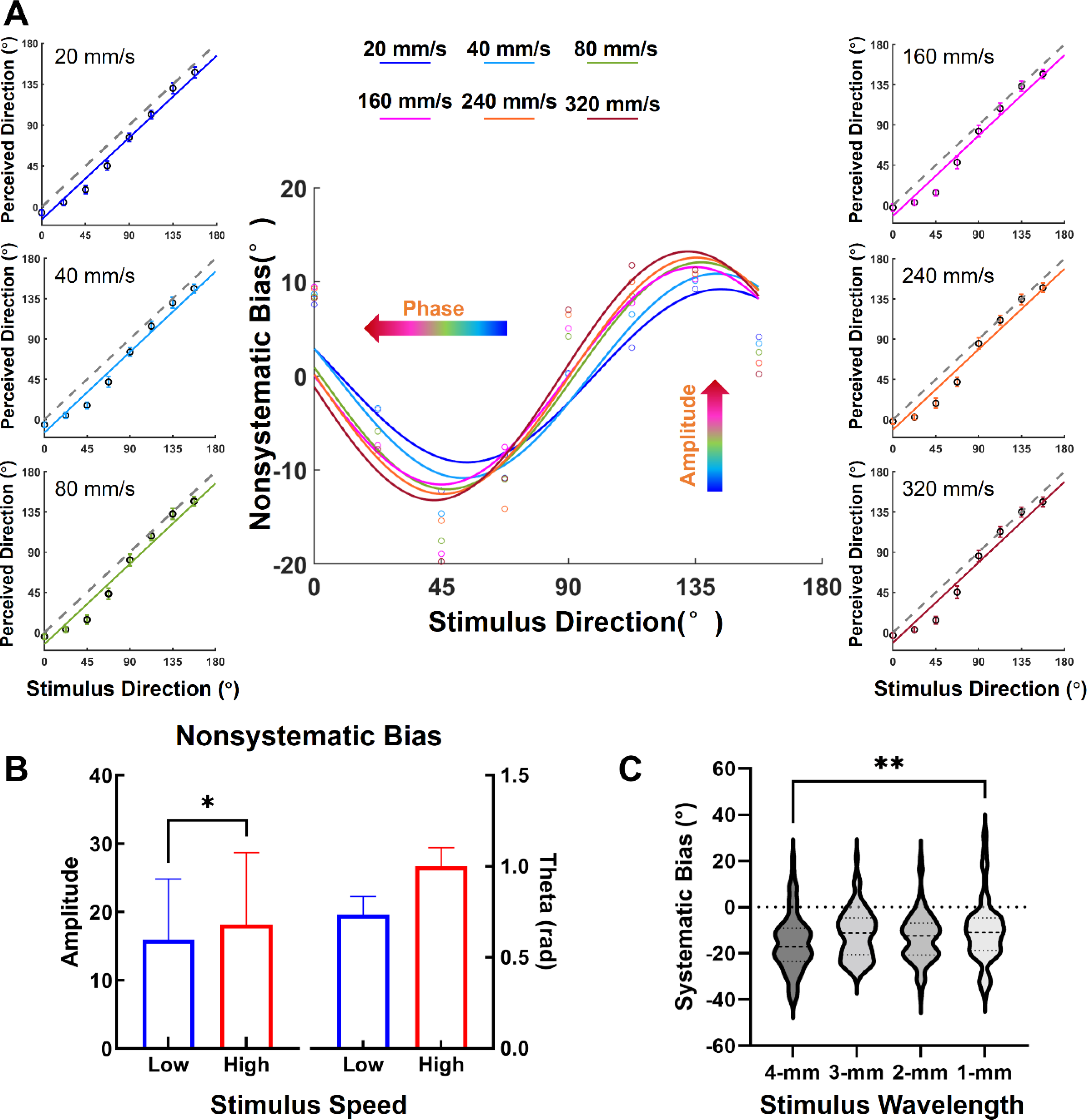
Direction bias for different tactile-motion stimulus speeds. **(A)** Perceived direction (black circles) of tactile-motion for the variety stimulus speeds (20, 40, 80, 160, 240, 320 mm/s) presented in two side columns. Different colored solid lines represent linear fits to systematic bias for different stimulus speeds, while the black dashed line represents the axis of symmetry between perceived direction and stimulus direction. In the middle, nonsystematic bias is illustrated with color dots, each associated with different cosine fitting curves. The data is further divided into low-speed (20,40 and 80 mm/s stimulus speeds) and high-speed (160, 240 and 320 mm/s) subgroups. **(B)** The amplitude of nonsystematic bias shows a significant different between low-speed and high-speed subgroups (*t*-test, T = 2.252, *p* < 0.05), while there was only a trend for phase of the nonsystematic bias. **(C)** Systematic bias exhibits a significant difference between the 1-mm and 4-mm stimulus ball wavelengths (F_(3, 384)_ = 3.330, *p* < 0.05; Bonferroni, *p* < 0.01 for 1-mm wavelength ball compared with 4-mm wavelength ball). Data is presented as the mean ± the standard error of mean (SEM).

The peak amplitudes and phases of nonsystematic bias exhibited a slight increase with stimulus speed. To analyze the influence of stimulus speeds on tactile-motion direction perception, data were categorized into low-speed and high-speed groups based on a goodness-of-fit curve (details in Supplementary, Figure S1). The low-speed group consisted of data from stimulus speeds of 20 mm/s, 40 mm/s, and 80 mm/s, while the high-speed group comprised data from 160 mm/s, 240 mm/s, and 320 mm/s. The amplitude of nonsystematic bias showed significant difference between the two groups (High-speed Group = 18.21±0.76, Low-Speed Group = 15.98±0.64; T (382) = 2.252, p < 0.05), while the phase of nonsystematic bias only exhibited a trend (High-speed Group = 0.74±0.10, Low-speed Group = 1.00±0.10; T (382) = 1.864, p = 0.063), as shown in Figure 3B. In addition, a trend was observed in the systematic bias between speed groups (High-speed Group = -11.58±0.88, Low-speed Group = -13.84±0.83, T (382) = 1.901, p = 0.058, data not shown). As shown in Figure 3C, systematic bias was significantly affected by the wavelength of stimulus ball (Figure 3C, F (3, 384) = 3.236, p < 0.05), specially between 1-mm wavelength and 4-mm wavelength balls (Bonferroni, p < 0.01). However, ball wavelength did not significantly affect nonsystematic bias, including amplitude (F (3, 384) = 0.352, p = 0.788) and only showed a trend for phase (F (3, 384) = 2.365, p = 0.071). These findings suggest that temporal features, represented by scanning speed, play a role in influencing nonsystematic bias, while spatial features, represented by the wavelength of the stimulus ball, impact systematic bias. This indicates that both temporal and spatial features are significant contributors to the tactile-motion direction perception.

### Effect of spatial features and velocity on tactile-motion speed perception

We further investigated the impact of spatial features (ball wavelength) and on tactile-motion speed perception. Normalized perceived speed was significantly modulated by the stimulus ball wavelength (F (3,264) = 33.110, p < 0.001), as shown in Figure 4A. Specifically, 1-mm and 2-mm wavelength balls elicited higher perceived speeds compared to 3-mm and 4-mm wavelength balls (Bonferroni, p < 0.001 for 1-mm ball vs. 3-mm or 4-mm ball, p < 0.05 for 2-mm ball vs. 3-mm or 4-mm ball). The effects of wavelength reached a lower plateau with 3-mm and 4-mm wavelength balls showing almost identical curves (p = 1). This suggests that the existence of a spatial resolution threshold, with 3-mm wavelength ball potentially representing the upper limit of spatial features capable of impacting speed perception. In summary, for speeds above 20 mm/sec and spatial frequencies between 1- and 3-mm spatial features play a significant role in modulating tactile-motion speed perception, indicating the presence of a spatial resolution threshold.

**Fig. 4.**
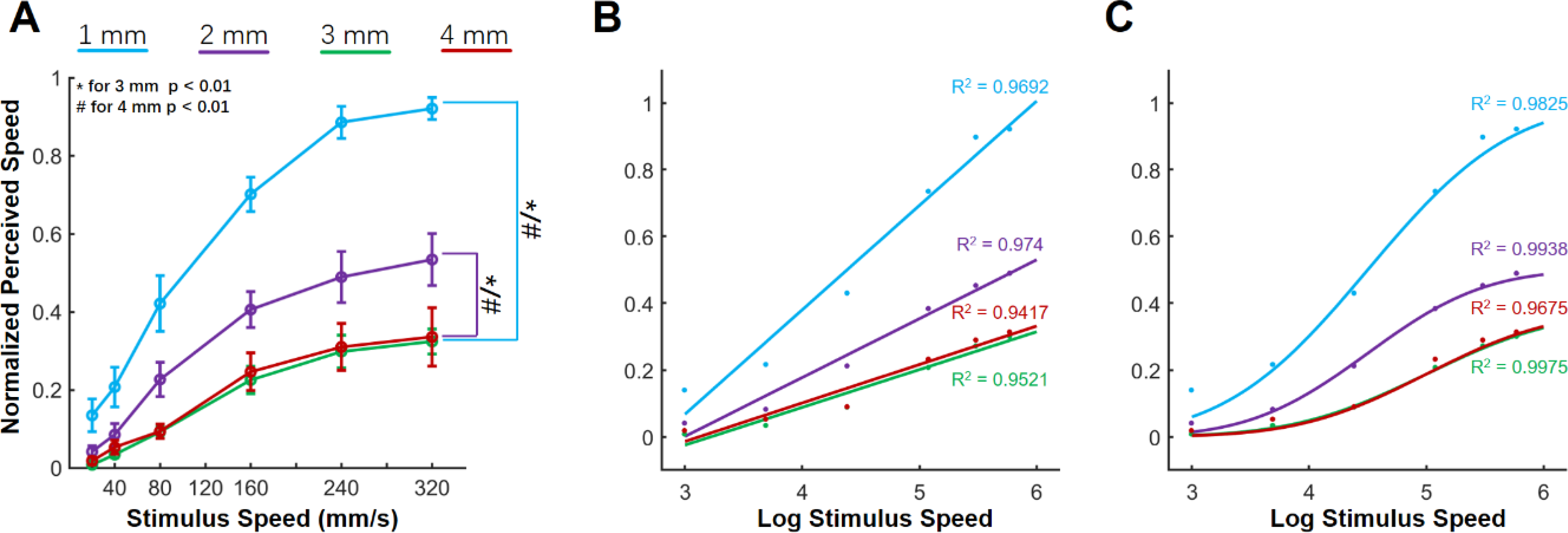
Normalized perceived speed for different stimulus ball wavelengths. **(A)** The plot shows the normalized perceived speeds as a function of stimulus speed. It demonstrates an increase in perceived speed with increasing stimulus speed that increases with ball wavelength. Compared with the 1-mm and 2-mm ball wavelengths, the perceived speeds evoked by the 3-mm and 4-mm ball wavelengths were significantly lower. Data is presented as mean ± standard error of mean (SEM). Two fitting curves were conducted to analyze these data: **(B)** linear function, and **(C)** Gaussian distribution-based psychometric function.

We used both linear and nonlinear functions to model the relationship between perceived speed and stimulus velocity and wavelength. The linear model was used to assess whether there was a simple, monotonic increase in tactile-motion speed perception, as showed in Figure 4B. Despite achieving R-square values exceeding 0.94, the results of the linear fitting suggested a limited effect of spatial features on perceived speed, poorly captured due to an unequal distribution of the sum of squares due to the errors (SSE) (see Figure 4B). To gain a more comprehensive understanding of the role of spatial features in tactile-motion speed perception, a Gaussian distribution-based psychometric function was used as a nonlinear descriptive model (Equation 6, Supplementary Materials), with the results of the fit depicted in Figure 4C. This model provided an averaged R-square of 0.98, indicating a superior fit compared to the linear models (details in Supplementary, Table S1). The Gaussian fit revealed three aspects of the psychometric function: an approximately linear increase at lower speeds, heightened sensitivity to speed changes at intermediate speeds (around 90 mm/s), and saturation at high speeds. These results suggest that the Gaussian distribution-based psychometric function offers a more comprehensive computational framework for understanding tactile-motion speed perception, involving nonlinear mechanisms that extend beyond linear trends.

## Discussion

The present study aimed to unravel the intricate interplay of spatial-temporal information in tactile-motion perception. Our investigation uncovers directional symmetry in tactile motion processing, explores interactions between spatial and temporal features, and proposes a fitting function for tactile-motion speed perception, seeking to enhance our understanding of tactile motion processing. One of the key findings of our study is that the anisotropic distortion of perceptual bias is quadrant-dependent, rather than specific to a preferred orientation, showing direction motion symmetry. Furthermore, we discovered that spatial and temporal features have divergent influences on tactile-motion perception. Spatial features primarily affect systematic bias, while temporal features exert their influence on nonsystematic bias. Moreover, our results revealed that the processing of tactile-motion speed encompasses both linear and nonlinear components, each associated with specific speed ranges. Collectively, these findings underscore the distinct roles played by spatial and temporal features in the integration of information crucial for tactile-motion perception.

Anisotropy in tactile perception of fingertip orientation is well-established, linked to variations in discrimination performance associated with the heterogeneous distribution of skin mechanoreceptors (*29*). Our study expanded this understanding in tactile direction perception with directions of motion covering a full 360 degrees, we observed a significant increase in perceptual biases in the first quadrant (Q1, upper right) and third quadrant (Q3, lower left), with more veridical performance in Q2 (upper left) and Q4 (lower right). This orthogonal diagonal pattern across the distal fingerpad, with greater sensibility along the Q2-Q4 axis and reduced sensibility along the Q1-Q3 axis. We propose this direction-specific perception along the Q2-Q4 axis can be attributed to natural hand placement posture and tactile exploration. For example, when the hand rests naturally on the tabletop, the arm forms an angle of approximately 45 degrees with the body-centered coordinate system, positioning the Q2-Q4 axis on the fingertip of the left index finger as a horizontal axis centered around the body. Furthermore, when exploring the texture of an object with the hand, movement along the Q2-Q4 axis on the fingertip of the left index finger is preferred over the Q1-Q3 axis. Understanding this directional sensibility sheds light on the interaction between mechanoreceptor distribution, neural processing, and hand posture in shaping perceptual biases.

A novel observation in our study was polar angle asymmetry in tactile perceptual performance when stimuli encompass the entire fingerpad. This differs from visual polar angle asymmetry (*30-33*), where visual anisotropy is generally better along the horizontal axis and the lower vertical axis (*34*). Visual polar angle asymmetries are influenced by factors such as photoreceptor density, retinal ganglion cells (*35, 36*), and cortical tissue distribution (*37-40*) in both humans and monkeys. In contrast, research on perceptual asymmetry in tactile studies is limited, often assumed to be related to the distribution of receptors and their corresponding neurons in somatosensory cortex. These polar angle asymmetries suggested inherent limitations in both tactile and visual system regarding location-specific sampling, restricting the amount of information transmitted from peripheral receptors to the brain. Furthermore, we observed that polar angle asymmetries may exhibit a unique form of symmetry in the Cartesian coordinate system. We categorized the perceptual biases based on various symmetry in Cartesian coordinate system. Our findings reveal that tactile perceptual performance exhibited central symmetry during direction discrimination, with neither vertical symmetry (X-axis) nor horizontal symmetry (Y-axis) playing the same role in the process (Figure 2C). In contrast, visual perception exhibits Y-axis symmetry (*41*). The difference between tactile and visual perceptions may be attributed to variations in neural circuits and computational processing models. In summary, our novel observation of quadrant-dependent anisotropic distortion with central symmetry enriches our understanding of tactile perceptual asymmetry.

To comprehend spatiotemporal interactions, we categorized perceptual bias into systematic and nonsystematic biases, which was proposed by Pei et al. (*42*). Systematic bias serves as a reliable indicator of the relative posture between different body parts, reflecting multisensory in integration. Interestingly, we observed a significant difference in systematic bias between coarse (4-mm ball) and dense (1-mm ball) wavelengths (Figure 3C), while stimulus speed did not affect it, consistent with Kuroki and Nishidazozi findings (*43*). This suggests that spatial features, rather than temporal features, influence systematic bias, suggesting a spatiotemporal segregation mechanism in tactile perception. Nonsystematic bias retains dynamic information, particularly temporal features. Although its origin likely arises from neural directional preferences or receptor responses, our findings reveal specific effects on these distinct properties during of tactile-motion perception. Specifically, the amplitude of the nonsystematic bias varied significantly among stimulus speed subgroups (Figure 3B), while the phase of the nonsystematic bias showed a subtle, nonsignificant shift due to speed. These findings suggest that temporal features can distort direction perception. Previous research indicates that faster arm movements lead to larger localization errors, linking temporal features to spatial errors in dynamic tactile perception (*44*). A study by Pei and associates (*42*) showed changes in hand-head position mainly influence systematic bias, leaving nonsystematic bias unaffected. Collectively, these and our findings suggest that temporal features play a vital role in the spatiotemporal integration of tactile perception, particularly regarding the alterations in direction perception.

Tactile speed perception is a complex process dependent on the interplay of spatial and temporal stimulus features, underpinned by neural mechanisms (*25, 26, 45*). Our study demonstrated that perceived speed depends on the spatial feature (ball wavelength) and stimulus speed (Figure 4A). Denser wavelengths increase the range of perceived speeds, suggesting precise textures could facilitate easier feature tracking, since denser wavelengths provide more detailed information about the surface texture, allowing for more accurate tracking of changes in motion. However, there is a spatial limit for this influence, as 3-mm and 4-mm ball wavelengths lead to similar speed perception, potentially related to the threshold for two-point discrimination on the fingerpad (*46*). Notably, smooth-surfaced stimuli result in increased speed discrimination errors (*47*). Furthermore, tactile speed perception is not solely dictated by peripheral mechanoreceptor distribution, as different afferents, particularly PC afferents, exhibit preferences for specific spatial frequency ranges (*24*). These findings reveal the intricate nature of tactile speed perception and its dependence on various sensory elements.

Previous research has established a monotonic relationship between perceived speed and stimulus speed under various spatial features (*25, 48*). Our data (Figure 4A) demonstrates these findings for stimulus speeds below 160 mm/s. However, at higher stimulus speed, the perceived speed plateaus across all ball wavelengths. While a linear function provides a good fit based on R-square values, the significant dispersion in SSE suggests the linear fits do not capture completely the underlying relationship. This implies the linear function is insufficient for a broader speed perception range. Tactile speed perception is believed to involve nonlinear neural coding resulting from the interaction of various types of mechanoreceptors relaying different spatial-temporal information (*49*). Although temporal features contribute to speed perception in both tactile and visual processing, tactile speed perception differs significantly from vision. Visual speed perception involves multiple processing stages within the visual pathway and is relatively well understood (*50-52*). In contrast, tactile speed perception may be influenced by the sparse innervation density and longer delays in mediating tactile speed information (*53*).

Given the spatial and temporal interactions in tactile speed perception, we introduced a Gaussian distribution-based psychometric function to incorporate both spatial and temporal features into a computational model (details in Supplementary, Table S1, Figure 4C). This function proposes a nonlinear integration of spatial and temporal information to better explain tactile speed perception, resulting in improved goodness-of-fit values for both R-square and SSE. This approach differs from a previous study by Greenspon et.al (*26*), which identified nonlinear relations in the power spectral densities of vibrations and speed, suggesting an interaction between spatial and temporal features in speed perception. Our findings (Table S1) indicate that the proposed function performs better with high spatial features (with 1-mm and 2-mm wavelength balls) compared to low spatial features (3-mm and 4-mm wavelength balls). The reduced range of perceived speeds is attributed to lower the density of elicited vibration by larger wavelengths is lower, resulting in reduced perceived speed (*25*). This suggests that coarse surfaces may have limitations in discriminating stimulus speed within a specific range.

The diversity of textures and shapes in the real-world results in high-dimension tactile sensations, and several studies have explored the role of motion in dynamic touch for tactile perception. Although our present study contributes valuable insights into the relationship between spatial and temporal features in tactile-motion perception, certain limitations should be acknowledged. For example, we used a simplified parameter “ball wavelength” to represent how stimulus texture interacts with tactile-motion processing. Furthermore, our study used a rotating tactile stimulus on a stationary finger to examine the integration of spatial and temporal features. Although these methods may not fully capture the complexities of real-world tactile experiences, our findings are intriguing and merit further investigations into their neural underpinnings. Future research endeavors that incorporate more natural textures and modes of tactile interaction may shed additional light on the intricate mechanisms of tactile-motion perception.

In conclusion, our study advances the understanding of tactile-motion processing by revealing quadrant-dependent anisotropic distortion, central symmetry, and the distinct roles of spatial and temporal features in shaping perceptual biases. The proposed Gaussian distribution-based psychometric function provides a nuanced framework for comprehending the nonlinear integration of spatial and temporal information in tactile speed perception. Further investigations into these mechanisms, considering the complexities of real-world tactile interactions, will contribute to a more comprehensive understanding of tactile-motion perception

## Materials and Methods

### Subject Details

This study was conducted with the approval of the Ethics Committee of the Second Affiliated Hospital of Zhejiang University School of Medicine (Approval No. 2015-081), and written informed consent was obtained from all participants. Thirty-three individuals were recruited and randomly divided into two groups. One for a tactile-motion direction symmetry experiment (n=17, mean age: 22.13±2.75 years, seven females) and the other for a tactile-motion speed integration experiment (n=16, mean age: 26.18±1.94 years, seven females). All participants were right-handed, as determined by the Edinburgh Handedness Inventory.

## Methods Details

### Stimuli Apparatus and Procedure

Participants were seated with their left forearm comfortably resting on holders, as shown in **Figure 1A**. The left forearm was vertically aligned with the elbow set at 90° angle, and the wrist, hand, palm up, and index finger (D2) were secured in a consistent posture to facilitate tactile stimulation. Participants’ head, eyes, and index finger were aligned along the posterior-to-anterior axis. A touchpad (PTH-660, Wacom Com Ltd., Japan) was placed between the participants and the tactile stimulator. Participants used their right index finger to draw an oriented line on the touchpad to indicate the perceived direction and speed of tactile stimuli. To mask any noise generated by the tactile stimulator, participants worn headphones emitting while noise.

Tactile stimuli were presented using a lab-designed ball stimulator, as described in (*54*). This stimulator provided three degrees of freedom, using one stepper and two servo motors to control the direction and speed of stimulation, as well as the relative distance between the stimulus ball and the distal fingerpad. Each stimulus ball had a diameter of 20 mm (**Figure 1B**) and featured four distinct wavelengths of square-wave gratings, each with the depth set to 500 μm and a duty cycle of 45%. To ensure a continuous perception of tactile-motion, the speed and direction of a stimulus were initiated before contact. The ball was then lowered at a speed of 5 mm/sec to achieve a 0.5-mm indentation, and stimulation lasted for 1 sec before the ball was lifted off from the finger. Before an experiment, the stimulus ball contacted the fingerpad to determine the starting position. The direction of motion was delineated within a planer coordinate system, with the upward direction (proximal to distal) along the index finger defined as 90°, as shown in **Figure 1C**.

Participants were instructed to respond by drawing oriented line on a touchpad to indicate direction and speed, reported by the length of the line. A point at the center of the touchpad marked the initial position of the reporting trajectory, and a circular region with 5-cm radius around the center served as a boundary. Following each stimulus trial, an auditory cue (a beep) prompted participants to report their tactile-motion perception. The trajectory of the drawn oriented lines was recorded using the Psychophysics Toolbox Version 3 (PTB-3, http://psychtoolbox.org) in MATLAB (R2018b, MathWorks, Natick, MA) at a sampling rate of 200 Hz. The perceived speed of the stimuli was calculated based on the number of pixels involved in the trajectories.

### Experimental Design

This study was designed to investigate the tactile directional symmetry of perceptual bias, and the influence of spatiotemporal features on tactile speed perception. To this end, two experiments were conducted as below.

### Experimental I: Tactile-motion directional symmetry

The primary objective of this experiment was to examine perceptual bias symmetry of the distal fingerpad in the perception of the tactile-motion direction and assess the impact of speed on this bias. A stimulus ball with a 4-mm wavelength was used to present stimuli in 24 directions, ranging from 0° to 360° in 15° increments, at stimulus speeds of 40 mm/s and 240 mm/s, as shown in **Figure 1D**. Each specific combination of stimulus parameters was presented five times, resulting in a total of 240 trials (24 directions × 2 speeds × 5 repetitions). To ensure reliability, the trials were divided into five blocks, each corresponding to a repetition of the stimuli. Participants had the flexibility to take breaks between these blocks, and the entire session took approximately 40 mins. Stimuli were presented in a pseudo-random order.

### Experimental II: Tactile-motion speed integration

Expanding on the insights gained from *Experiment I*, *Experiment II* introduced covarying spatial-temporal features to investigate their influence on tactile-motion speed perception. Four stimulus balls with four different wavelengths (1, 2, 3, and 4 mm), six distinct speeds (20, 40, 80, 160, 240 and 320 mm/s), and eight directions ranging from 0 to 157.5° with 22.5° increments were employed, as shown in **Figure 1D**. This comprehensive experiment included 960 trials (8 directions x 6 speeds x 5 repetitions x 4 wavelengths) and took about 180 minutes to complete. Participants had the flexibility to completed these sets over two days, with a 3-minute rest interval after every 48 trials (8 directions x 6 speeds) to prevent attention fatigue.

## Data Processing

During each trial, the positional time series of the oriented line on the touchpad was recorded. This recorded signal was differentiated to calculate the perceived speed, while direction was calculated using the arctangent function, tan^−1^(Y/X), for the entire trajectory. In addition, an average perceived direction and speed was computed based on the five stimulus repetitions.

Perceptual bias *B*_*d*_ was defined as the difference between the perceived direction *D*_*p*_ and the stimulus direction *D*_*s*_:

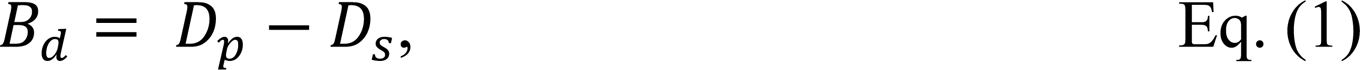

A negative *B*_*d*_ implies a clockwise shift in direction perception, indicating that participants tend to perceive the direction as being rotated clockwise compared to its stimulus direction. Conversely, a positive *B*_*d*_ points to a counterclockwise shift in direction perception.

Perceptual bias was separated into systematic bias and nonsystematic bias, representing inherent and stimulus specific responses, respectively. The systematic bias *B*_*s*_ was defined as the mean of the perceptual bias *B*_*d*_ :

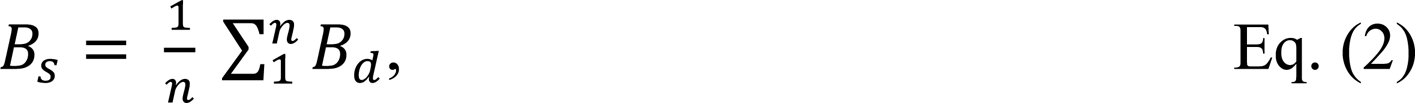

where *n* was the number of stimulus directions. The nonsystematic bias, *B*_*ns*_, calculated the difference of the perceptual bias for each motion direction from the overall perceptual bias (*42*). The calculation can be expressed as follows:

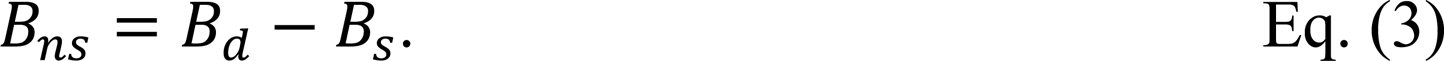

In our study, a cosine function was employed as the fitting curve:

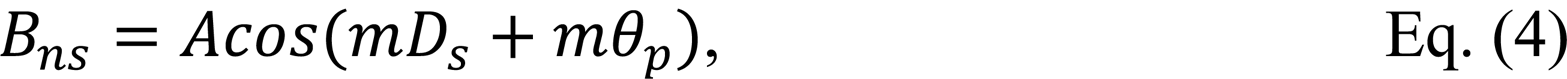

where *m* was set to 2, as in a previous study (*42*), to ensure a best fitting curve for our dataset. Parameters *A* and *θ*_*p*_ represent the peak amplitude and peak phase of the nonsystematic bias, respectively.

To analyze tactile-motion speed perception, perceived speed was standardized using Z-score normalization, and both linear and nonlinear models were used to explore interactions between stimuli and perceived speed. For linear fitting, the equation adopted was as follows:

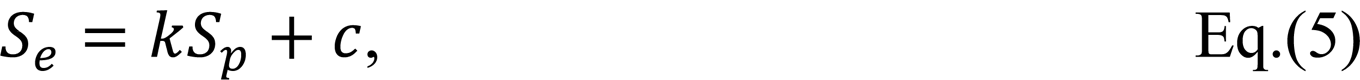

where *S_e_* was the estimated normalized perceived speed, *S*_*p*_ corresponded to the normalized perceived speed based on a z-score for each participant across the entire experiment, and *k* denoted the slope and *c* denoted the intercept of the linear model.

To incorporate presumed mechanisms underlying the tactile-motion perception process, a Gaussian distribution-based psychometric function was used to fit the data:

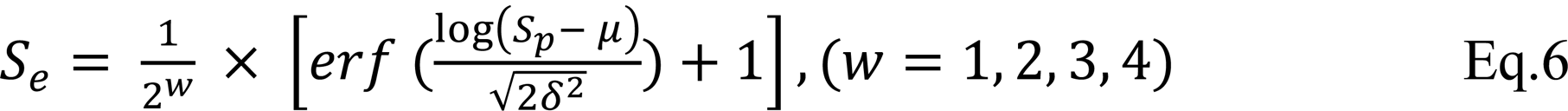

where *w* was the wavelength of the stimulus ball, *μ* was the center threshold of perceived speed, and *δ* was a constant corresponding to the just-noticeable difference. We employed the Gaussian error function (*erf*) available in MATLAB to analyze the data and model specific aspects of the observed phenomena. This model postulates that stimulus wavelength is a predominant factor influencing speed perception, and *μ* was set as 90 mm/s based on the value associated with the highest R-square (details in Supplementary, **Figure S1**). Both linear and nonlinear models were implemented using *Curve Fitting Toolbox in* MATLAB (2021a) for regression analysis.

## Statistical Analysis

All the statistical analyses were conducted using SPSS software (version 22, IBM Corp., Armonk, NY). Normality was assessed using the Shapiro-Wilk test and Levene test. A Student’s *t* test was used to evaluated the influence of stimulus speed on mean perceptual bias across stimulus directions. Repeated-measure ANOVA was conducted for each quadrant to investigate potential patterns of anisotropic distortion across stimulus directions. Pearson’s correlation was applied to assess potential symmetry, including X-axial, Y-axial and Central-axial symmetries. Repeated-measures ANOVA was used to assess the impact stimulus speed on direction perception, including systematic bias, amplitude of nonsystematic bias, and phase of nonsystematic bias. ANOVAs were also performed to assess the effects of stimulus wavelengths on tactile direction perception. To explore the effects of wavelength on tactile speed perception, ANOVA was first used to examine the difference as the stimulus speed varied, followed by a Pearson’s correlation to assess the consistency between the two curves (3-mm and 4-mm).

## Acknowledgments

We are grateful for the valuable feedback provided by Bensmaia J. Sliman and Anna W. Roe. Special thanks to Yu-Cheng Pei and Mengjie Xin for their assistance in the initial stages of this research.

## Funding

STI 2030—Major Projects 2021ZD0200401 (HYL, GP, KES)

National Key R&D Program of China 2021YFF0702200 (HYL)

National Natural Science Foundation of China 82101323 (HYL)

Key R&D Program of Zhejiang Province 2021C03001 (HYL)

Fundamental Research Funds for the Central Universities 2019XZZX003-20 (HYL)

## Author contributions

Conceptualization: HYL, BQ, RMF

Methodology: BQ, XT, GP

Investigation: BQ, XT, ZT, HW

Visualization: BQ, LL

Supervision: HYL

Writing—original draft: BQ, KES, HYL

Writing—review & editing: HYL, RMF, BQ

## Data and materials availability

All data are available in the main text or the supplementary materials.

