## Supplemental Figure 1 and Supplemental Table 1 for "Unveiling interactions of spatial-temporal information in tactile motion perception"

Science Advances
AAAS

### Supplementary Materials for

#### Unveiling interactions of spatial-temporal information in tactile motion perception

Boyi Qu *et al.*

**This PDF file includes:**

Supplementary Text
Figs. S1
Tables S1

**Supplementary Text**

*Model fit performance all the wavelengths and stimulus speeds.*

To investigate the potential role of temporal and spatial features in tactile-motion speed perception, we introduced a Gaussian distribution-based psychometric function as following:

$$S_e = \frac{1}{2^w} \times \left[\mathrm{erf}\left(\frac{\log(S_p - \mu)}{\sqrt{2\delta^2}} + 1\right], (w = 1, 2, 3, 4) \qquad \text{Eq. S1}$$

Here, $S_p$ is the normalized perceived speed, $S_e$ is estimated normalized perceived speed, $w$ is the wavelength of stimulus ball, $\mu$ is the center threshold of perceived speed and $\delta$ is a free parameter. To optimize the performance of the fitting function, we determined a center threshold based on a goodness-of-fit test. We considered speeds within the range of 60 mm/s to 240 mm/s, with the step by 0.1 logarithmic stimulus speed as the potential center threshold values, and evaluated the performance for each wavelength. The goodness-of-fit test was conducted using the MATLAB R2021b Curve Fitting Toolbox. As shown in Figure S1, the $R^2$ values for the 1-mm, 2-mm wavelengths peak at 4.5 logarithmic stimulus speed (corresponding to a stimulus speed of 90 mm/s, marked by the dashed red line). Conversely, the 3-mm and 4-mm wavelengths peaked at a 4.8 logarithmic stimulus speed (corresponding to a stimulus speed of 120 mm/s, marked by a dashed black line). For all types of balls, we defined the center threshold $\mu$ based on the peak values in our current study. To assess the potential impact of stimulus speeds on tactile-motion direction perception, we divided the data into low-speed (20 mm/s, 40 mm/s, and 80 mm/s) and high-speed groups (160 mm/s, 240 mm/s, and 320 mm/s) based on the goodness-of-fit curve.

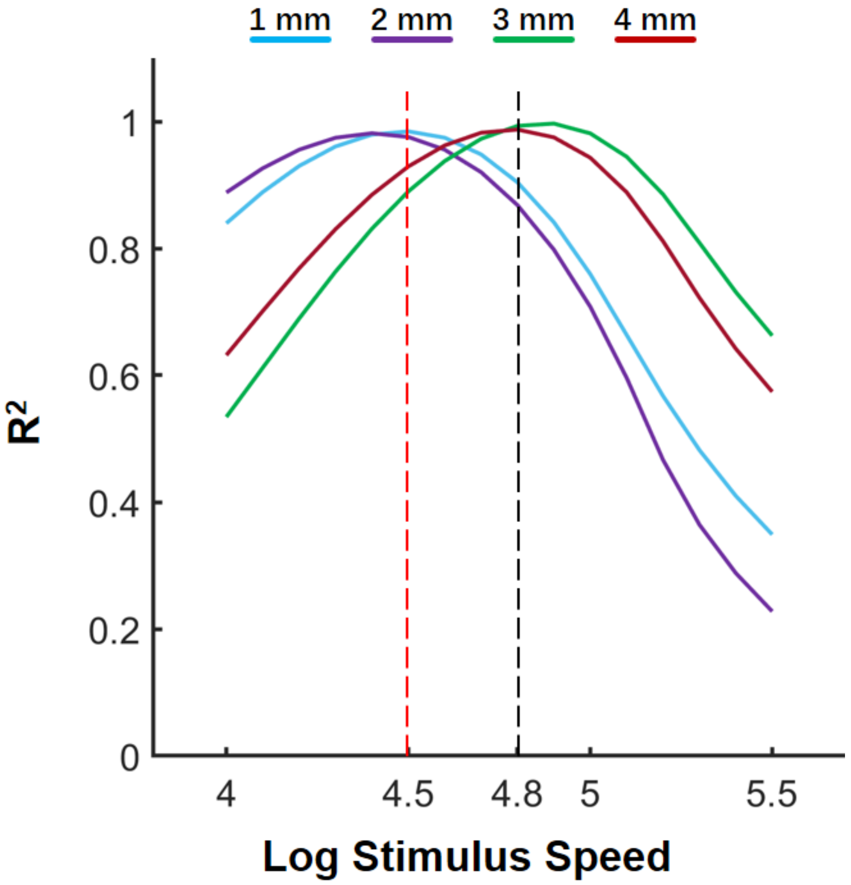


**Fig. S1. Model fit performance all the wavelengths and stimulus speeds.** The goodness-of-fit peaks showed at logarithmic stimulus speed of 4.5 for the 1-mm and 2-mm wavelength, and logarithmic stimulus speed of 4.8 for the 3-mm and 4-mm wavelengths (dashed black line). The peaks in goodness of fits were used as center thresholds ($\mu$) in Eq. S1 and manuscript Eq. (6).

**Table S1.**

| The sum of squares due to error of fitting curves | | |
|---|---|---|
| **Wavelength /mm** | **linear fitting function** | **nonlinear fitting function** |
| **1** | 0.01805 | 0.01025 |
| **2** | 0.004831 | 0.001143 |
| **3** | 0.003741 | 0.0001916 |
| **4** | 0.004772 | 0.002658 |

The Gaussian based psychometric function produced better fits than the linear model for all ball wavelengths.
